# CD155 blockade enhances allogeneic natural killer cell-mediated antitumor response against osteosarcoma

**DOI:** 10.1101/2023.06.07.544144

**Authors:** Monica M Cho, Longzhen Song, Aicha E Quamine, Fernanda Szewc, Lei Shi, Johnathan D Ebben, David P Turicek, Jillian M Kline, Devin M Burpee, Emily O Lafeber, Madison F Phillips, Amanda S Ceas, Amy K Erbe, Christian M Capitini

## Abstract

**Background:** Allogeneic bone marrow transplant (alloBMT) is curative for hematologic malignancies through the graft-versus-tumor (GVT) effect but has been ineffective for solid tumors like osteosarcoma (OS). OS expresses CD155 which interacts strongly with inhibitory receptors TIGIT and CD96 but also binds to activating receptor DNAM-1 on natural killer (NK) cells. CD155 has never been targeted after alloBMT. Combining adoptively transferred allogeneic NK (alloNK) cells with CD155 blockade after alloBMT may enhance a GVT effect against OS.

**Methods:** Murine NK cells were activated and expanded ex vivo with soluble IL-15/IL-15Rα. AlloNK and syngeneic NK (synNK) cell phenotype, cytotoxicity, cytokine production, and degranulation against CD155-expressing murine OS cell line K7M2 were assessed in vitro. Mice bearing pulmonary OS metastases underwent alloBMT and alloNK cell infusion with anti-CD155 either before or after tumor induction, with select groups receiving anti-DNAM-1 pretreated alloNK cells. Tumor growth, GVHD and survival were monitored, and differential gene expression of lung tissue was assessed by RNA microarray.

**Results:** AlloNK cells exhibited superior cytotoxicity against CD155-expressing OS compared to synNK cells, and this activity was enhanced by CD155 blockade. CD155 blockade increased alloNK cell degranulation and interferon gamma production through DNAM-1. In vivo, CD155 blockade with alloNK infusion increased survival when treating OS that relapsed after alloBMT. No benefit was seen for treating established OS before alloBMT. Treatment with combination CD155 and anti-DNAM-1 pretreated alloNK ameliorated survival and tumor control benefits seen with CD155 blockade alone. RNA microarray showed mice treated with alloNK and CD155 blockade had increased expression of cytotoxicity genes and the NKG2D ligand H60a, whereas mice treated with anti-DNAM-1 pretreated alloNK cells resulted in upregulation of NK cell inhibitory receptor genes. Whereas blocking DNAM-1 on alloNK abrogated cytotoxicity, blocking NKG2D had no effect, implying DNAM-1:CD155 engagement drives alloNK activation against OS.

**Conclusions:** These results demonstrate the safety and efficacy of infusing alloNK cells with CD155 blockade to mount a GVT effect against OS and show benefits are in part through DNAM-1. Defining the hierarchy of receptors that govern alloNK responses is critical to translating alloNK cell infusions and immune checkpoint inhibition for solid tumors treated with alloBMT.

**WHAT IS ALREADY KNOWN ON THIS TOPIC:**

- Allogeneic bone marrow transplant (alloBMT) has yet to show efficacy in treating solid tumors, such as osteosarcoma (OS). CD155 is expressed on OS and interacts with natural killer (NK) cell receptors, such as activating receptor DNAM-1 and inhibitory receptors TIGIT and CD96 and has a dominant inhibitory effect on NK cell activity. Targeting CD155 interactions on allogeneic NK cells could enhance anti-OS responses, but this has not been tested after alloBMT.

**WHAT THIS STUDY ADDS:**

- CD155 blockade enhances allogeneic natural killer cell-mediated cytotoxicity against OS and improved event-free survival after alloBMT in an in vivo mouse model of metastatic pulmonary OS. Addition of DNAM-1 blockade abrogated CD155 blockade-enhanced allogeneic NK cell antitumor responses.

**HOW THIS STUDY MIGHT AFFECT RESEARCH, PRACTICE OR POLICY:**

- These results demonstrate efficacy of allogeneic NK cells combined with CD155 blockade to mount an antitumor response against CD155-expressing OS. Translation of combination adoptive NK cell and CD155 axis modulation offers a platform for alloBMT treatment approaches for pediatric patients with relapsed and refractory solid tumors.

## INTRODUCTION

Osteosarcoma (OS) is the most common primary bone cancer in children, with approximately 1,000 cases per year in the U.S. While patients with localized disease have a 5-year event free survival (EFS) rate of 70%, patients with metastatic disease have only a 24% survival rate (1,2). There are no curative treatment options for patients with metastatic or relapsed/refractory disease.

Natural killer (NK) cells are innate lymphocytes that are cytotoxic against tumors based on the balance between activating and inhibitory signals from cell surface receptor engagement. NK cell self-tolerance prevents healthy tissue damage through interaction between inhibitory killer-immunoglobulin-like receptors (KIRs), analogous to the murine Ly49 family of receptors, and their ligands, self-major histocompatibility complex (MHC) class I molecules (3). Malignant cells not recognized as “self” due to KIR/Ly49 receptor mismatch are susceptible to lysis because these inhibitory receptors cannot engage MHC I ligand. Allogeneic hematopoietic stem cell transplant (alloHSCT) combined with adoptively transferred allogeneic NK cells is an attractive approach to augment the graft-versus-tumor (GVT) effect generated by KIR mismatch (4). Graft-versus-host-disease (GVHD) is T-cell mediated and a major barrier in applying alloHSCT for solid tumors; thus, T-cell depletion of donor grafts may reduce the risk of GVHD while administering other effector cells like NK cells post-HSCT (5,6). While several studies observe benefit of KIR-mismatched NK cells in leukemias, more studies are needed to demonstrate KIR mismatch benefits in pediatric solid tumors (7–11).

Although allogeneic NK (alloNK) cells have potential to treat solid tumors, there are significant barriers with respect to *in vivo* function. AlloNK cells with a KIR/KIR-ligand mismatch have not shown potent initial antitumor response due to limited function and persistence. To overcome these barriers, ex vivo activated and expanded (EAE) NK cells can boost antitumor effects (12). Compared to resting NK cells, EAE NK cells use multiple synergistic activating receptors and exhibit superior target cell lysis (13). EAE NK cells more effectively lyse OS cell lines and patient-derived chemotherapy-resistant OS compared to resting NK cells (14). Since IL-15 levels increase more than 10-fold during conditioning prior to HSCT peaking at 15 days post-HSCT, alloHSCT is an excellent platform for EAE alloNK cell therapy, since IL-15 supports NK cell engraftment, activation, and persistence. Since adoptively transferred NK cells can match the HSCT donor graft, which itself generates NK cells early post-HSCT, there is no risk of rejection (15,16).

Overcoming tumor microenvironment (TME) immunosuppression is a major barrier to successful adoptive NK cell therapy. While many activating receptors on NK cells have been implicated in driving the GVT effect (e.g. KIR, NKG2D, NKG2C, etc), their relative hierarchy is still unknown. CD155, originally discovered as the poliovirus receptor, is an immunoglobulin superfamily adhesion molecule involved in cell adhesion and migration, proliferation, and immune modulation (17). Acute myeloid leukemia, multiple myeloma, neuroblastoma, fibrosarcoma, glioblastoma, melanoma, prostate cancer, renal cell cancer, pancreatic carcinomas, colon, non-small cell lung, ovarian, and breast carcinomas, and OS have been found to express CD155 (18–21). High CD155 expression correlates with poor clinical outcomes and affects tumor susceptibility to NK cell lysis (22). DNAM-1 is a CD155 receptor critical for NK cell activation that synergizes with other activation pathways, increasing actin polymerization and granule polarization (23). DNAM-1 triggers direct NK cell cytotoxicity upon interaction with CD155 and Nectin2/CD112 (24). CD155 also interacts with inhibitory receptors, TIGIT and CD96. TIGIT is an immunoglobulin superfamily receptor expressed on T cells and NK cells. Upon CD155 and CD112 interaction, TIGIT attenuates NK cell degranulation, cytokine production, and cytotoxicity (25). TIGIT has higher affinity for CD155 and outcompetes DNAM-1, resulting in inhibitory signaling. CD96 also competes with DNAM-1 for CD155 binding and directly inhibits IFNγ production, activation, and cytotoxicity (26). Combining the effects of KIR/Ly49 mismatch and CD155 immune checkpoint blockade can potentially overcome TME immunosuppression. Prior studies suggest that CD155 blockade or knockout favors NK cell antitumor effects by preventing dominant inhibitory effects, but this has not been explored in the context of alloHSCT nor in combination with adoptively transferred EAE alloNK cells (21). We hypothesized that CD155 blockade after T cell depleted alloHSCT will enhance the GVT effect of adoptively transferred EAE alloNK cells by increasing NK activation and cytotoxicity without exacerbating GVHD.

## METHODS

### Mice

Female C57BL/6NCr (B6), B6-Ly5.1/Cr and BALB/cAnNCr (BALB/c) mice (Charles River Laboratories, Wilmington, MA) were housed in accordance with the Guide for Care and Use for Laboratory Mice. All animal experiments were performed under an IACUC approved protocol at the University of Wisconsin-Madison (M005915).

### Ex vivo activation and expansion of murine NK cells

Bone marrow was harvested from 8–12-week-old female B6 or BALB/c mice and processed into a single cell suspension and red blood cells were lysed with ACK lysis buffer (Lonza, Walkersville, MD). NK cells were isolated by magnetic cell separation with a NK cell isolation kit and an AutoMACS Pro (Miltenyi Biotec, San Jose, CA). NK cells were expanded and activated for 14 days using mouse 10 ng/ml IL-15/IL-15Rα conjugate, comprised of recombinant mouse IL-15 protein (R&D Systems, Minneapolis, MN) and recombinant mouse IL-15Ralpha Fc chimera protein (R&D Systems), in RPMI medium (Corning Life Sciences, Durham, NC) with 10% fetal bovine serum (GeminiBio, Sacramento, CA) with 50 IU/mL penicillin/streptomycin (Lonza), 0.11 mM 2-betamercaptoethanol (Gibco, Carlsbad, CA), 1X MEM nonessential amino acids (Corning), 25 mM HEPES buffer (Corning), and 1 mM sodium pyruvate (Corning). IL-15/IL-15 receptor conjugate was repleted during media changes with half fresh and half conditioned media every 2-3 days to maintain cell concentration of 1-2 × 10^6^ cells/mL. Over the 14-day expansion, freshly isolated and cultured live NK cells were counted and analyzed by flow cytometry at days 0, 7, 9, 12 and 14. NK cells were collected 12 days post-NK cell isolation for all other in vitro and in vivo assays.

### Bone marrow transplant in vivo model with adoptive NK cell transfer and antibody blockade treatment

Bone marrow cells were harvested from B6 mice and processed into single cell suspensions. A CD3ε microbead kit (Miltenyi Biotec) was used to deplete T cells from murine bone marrow cells. Recipient BALB/c mice were lethally irradiated with 8 Gy in two 4 Gy fractions using an X-Rad 320 (Precision Xray Irradiation, Madison, CT) and transplanted with 2.5-5E6 T cell-depleted B6 bone marrow cells by retroorbital injection. For an established OS disease alloHSCT model, on day 0, BALB/c mice were inoculated with 1E6 intravenous luciferase^+^ K7M2 murine OS cells. Tumor-bearing BALB/c mice underwent bone marrow transplant at day 7. For a post-HSCT relapsed disease model, BALB/c mice underwent alloBMT at day 0, followed by 1 x 10^6^ intravenous luciferase^+^ K7M2 murine OS cell tumor inoculation at day 7. For both models, 7-8 days after tumor injection, mice received 1 x 10^6^ allogeneic B6 EAE NK cells per mouse, expanded with IL-15/IL-15Rα for 12 days, pretreated with no antibody blockade, purified rat anti-mouse anti-DNAM-1 (Biolegend Cat. No. 128822, San Diego, CA), or purified rat anti-mouse anti-NKG2D (Biolegend Cat. No. 130202) for 30 minutes at 37°C and washed with PBS prior to retroorbital injection with concurrent intraperitoneal treatment by 200 ug of purified rat IgG2a antibody (Leinco Cat. No. I-1177, Fenton, MO) or purified rat anti-mouse anti-CD155 antibody (Leinco Cat. No. C2833).

### In vivo GVHD and tumor burden monitoring

Tumor burden in mice was monitored by bioluminescence using the IVIS Spectrum in vivo imaging system (Perkin-Elmer, Waltham, MA). Image data were analyzed using Living Image software (Perkin-Elmer, Waltham, MA). Treatment toxicity and GVHD were measured using body weight loss and GVHD clinical scoring system (27). EFS was calculated with an event defined as tumor burden increased by at least four-fold from initial tumor total flux (photons/s) or subject death.

### Statistical Analysis

Data presented show analyses of experimental replicate samples with plotted individual sample values, mean and standard error of the mean (SEM). Comparisons were performed using two-sided two-sample t-tests (two experimental groups) or one-way-ANOVA/ Kruskal-Wallis with Dunn’s multiple comparisons tests (greater than two experimental groups). Significance was determined by p-values ≤ 0.05. Statistical analyses were performed using Prism 9 (Graphpad Software, San Diego, CA).

For additional information, please refer to supplemental methods.

## RESULTS

### Expanded murine NK cells exhibit an activated phenotype

B6 and BALB/c NK cells were isolated from bone marrow and expanded with soluble IL-15/IL-15Rα complex (sIL-15/IL15Rα) for 14 days. NK cell yields per mouse post-isolation were 0.176 ± 0.021 × 10^6^ B6 NK cells/mouse and 0.568 ± 0.185 × 10^6^ BALB/c NK cells/mouse (Fig. 1A). Both B6 and BALB/c murine NK cells expanded considerably, and peak expansion was observed at day 12 with 15.92 ± 2.91 × 10^6^ B6 NK cells/mouse and 7.82 ± 2.54 × 10^6^ BALB/c NK cells/mouse (Fig. 1A). Maximal fold expansion occurred at day 12 with a fold expansion of 89.600 ± 13.728 for B6 NK cells and 16.988 ± 6.050 for BALB/c NK cells (Fig. 1B). At day 14, B6 and BALB/c NK cell counts fell to 2.128 ± 0.505 × 10^6^ B6 NK cells/mouse and 3.560 ± 6.07 × 10^6^ BALB/c NK cells/mouse (Fig. 1A).

**Figure 1.**
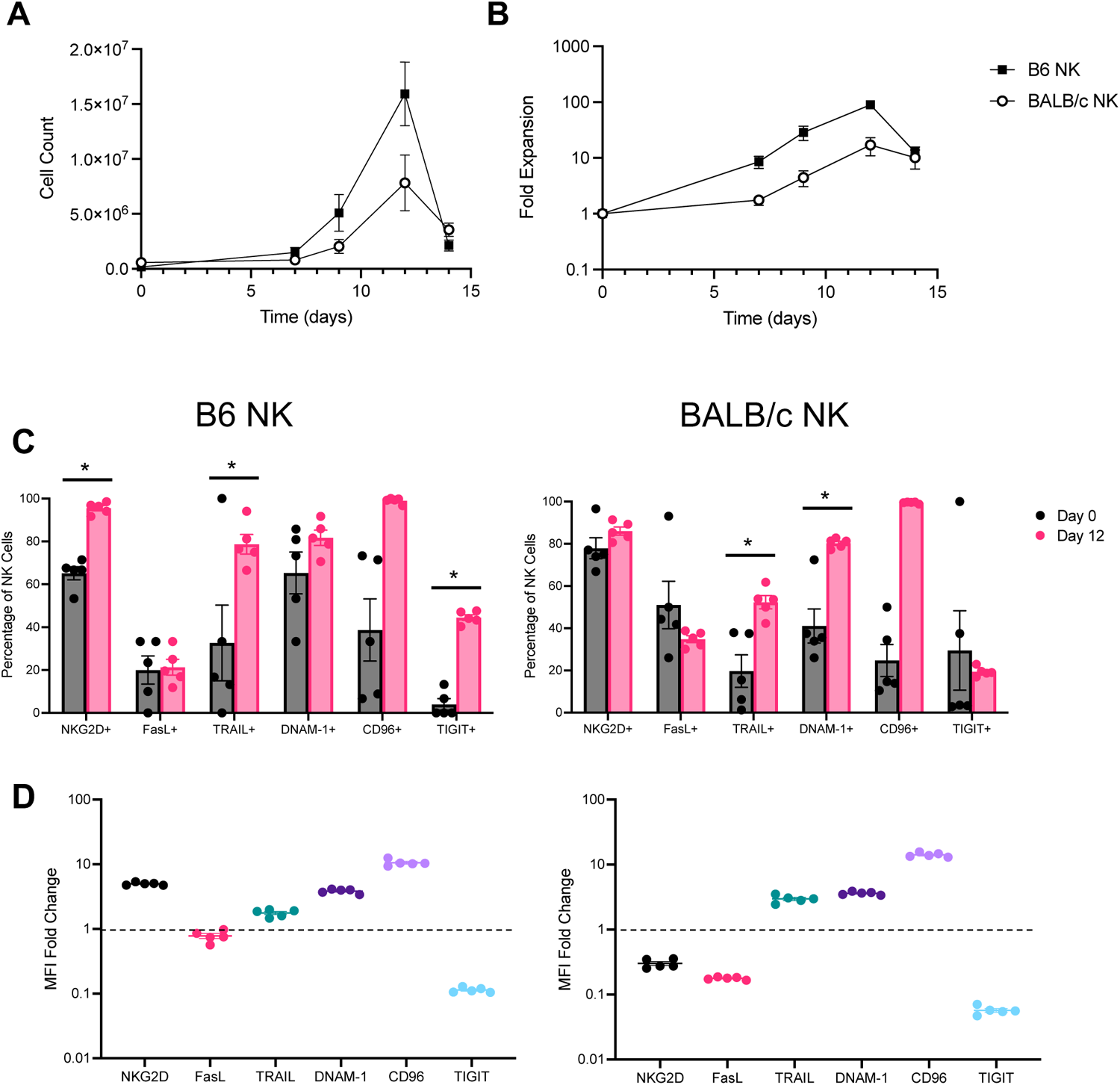
IL-15 expansion of murine NK cells results in activated murine NK. B6 and BALB/c NK cells cultured with IL-15/IL-15Rα conjugate with additional media and cytokine repletion every 2-3 days are shown. (A) Cell count and (B) fold expansion at days 0, 7. 9, 12, and 14. Percentage of (C) B6 and BALB/c NK cells with marker expression at day 0 and day 12 shown with individual experimental replicates plotted with mean and SEM. (D) Fold change of marker expression MFI at day 12 compared to day 0 shown for B6 NK cells and BALB/c NK cells. Mean with SEM shown for experimental replicates (n = 5). Two-sided two-sample t tests were performed (* p < 0.05, ** p < 0.01, *** p < 0.001, **** p > 0.0001).

B6 and BALB/c NK cell activation was assessed by flow cytometry. During expansion, the proportion of NKG2D^+^ cells increased, peaking at day 12, with 95.52 ± 2.654% NKG2D^+^ B6 NK cells and 86.02 ± 4.368% NKG2D^+^ BALB/c NK cells (Supplementary Fig. 1A-B). In B6 NK cells, %NKG2D^+^ B6 NK cells were significantly increased at day 12 compared to pre-expansion (Supplementary Fig. 1A). Thus, sIL-15/IL15Rα expands and activates NK cells from both mouse strains. To evaluate cytotoxic potential, tumor necrosis factor (TNF) family members FasL and TRAIL were measured. B6 and BALB/c NK cells did not exhibit significant increases in the median fluorescence intensity (MFI) of FasL or TRAIL, although the proportion of TRAIL^+^ B6 NK cells increased at days 12 and 14 (Supplementary Fig. 1A-B). BALB/c NK cells had an initial increase in TRAIL MFI and percentage of TRAIL^+^ NK cells at days 5 and 7, but by day 12, increased TRAIL MFI was the only persistent increase into the second week of expansion (Supplementary Fig. 1A-B).

The CD155 receptors DNAM-1, TIGIT and CD96 are expressed on mature NK cells, but it is unclear how sIL-15/IL15Rα affects their expression and if effects are strain specific. DNAM-1 MFI was significantly increased at day 12 in both B6 and BALB/c NK cells, while the proportion of DNAM-1^+^ NK cells was significantly increased in BALB/c NK cells (Supplementary Fig. 1C-D). CD96 MFI was significantly elevated in B6 and BALB/c NK cells (Supplementary Fig. 1C-D). TIGIT MFI and proportion of TIGIT^+^ NK cells were both significantly increased at day 12 of B6 NK cell expansion (Supplementary Fig. 1C-D). After 12 days of culture, there were increased proportions of B6 NK cells expressing NKG2D, TRAIL, and TIGIT, and there were greater proportions of BALB/c NK cells expressing TRAIL and DNAM-1 (Fig. 1C). With respect to MFI fold change at day 12, murine NK cells exhibited significant increases in NKG2D, TRAIL, DNAM-1 and CD96 expression (Fig. 1D). At day 14, there was marked loss of B6 and BALB/c NK cells, possibly due to increased exhaustion evidenced by increased LAG-3 and TIM-3 expression (Supplementary Fig. 2A-B). Most significant changes in NK phenotype and proliferation coalesced on day 12, so NK cells expanded for 12 days with sIL-15/IL15Rα were used for subsequent experiments.

### CD155 blockade enhances alloreactive NK cell activity against OS

To assess murine NK cell activity against OS, we used a CD155-expressing murine K7M2 OS cell line derived from a BALB/c mouse (Fig. 2A). K7M2 had a low proportion of CD112^+^ cells (Fig. 2A). K7M2 OS also expressed Fas and TRAIL receptor 2, which bind to apoptosis-inducing FasL and TRAIL on NK cells (Fig. 2B). To determine the contribution of Ly49 mismatch (where inhibitory Ly49 receptors on NK cells cannot recognize cognate MHC I ligand) to anti-OS activity, allogeneic NK cells from B6 mice (alloNKs) and control syngeneic Ly49-matched NK cells from BALB/c mice (synNKs) were co-cultured with K7M2 OS and evaluated for cytotoxicity using a live cell imaging system. Compared to synNKs, alloNKs showed significantly higher percentage of OS lysis after 24 hours (alloNK 59.13 ± 3.50%, synNK 30.13 ± 3.45%) (Fig. 2C).

**Figure 2:**
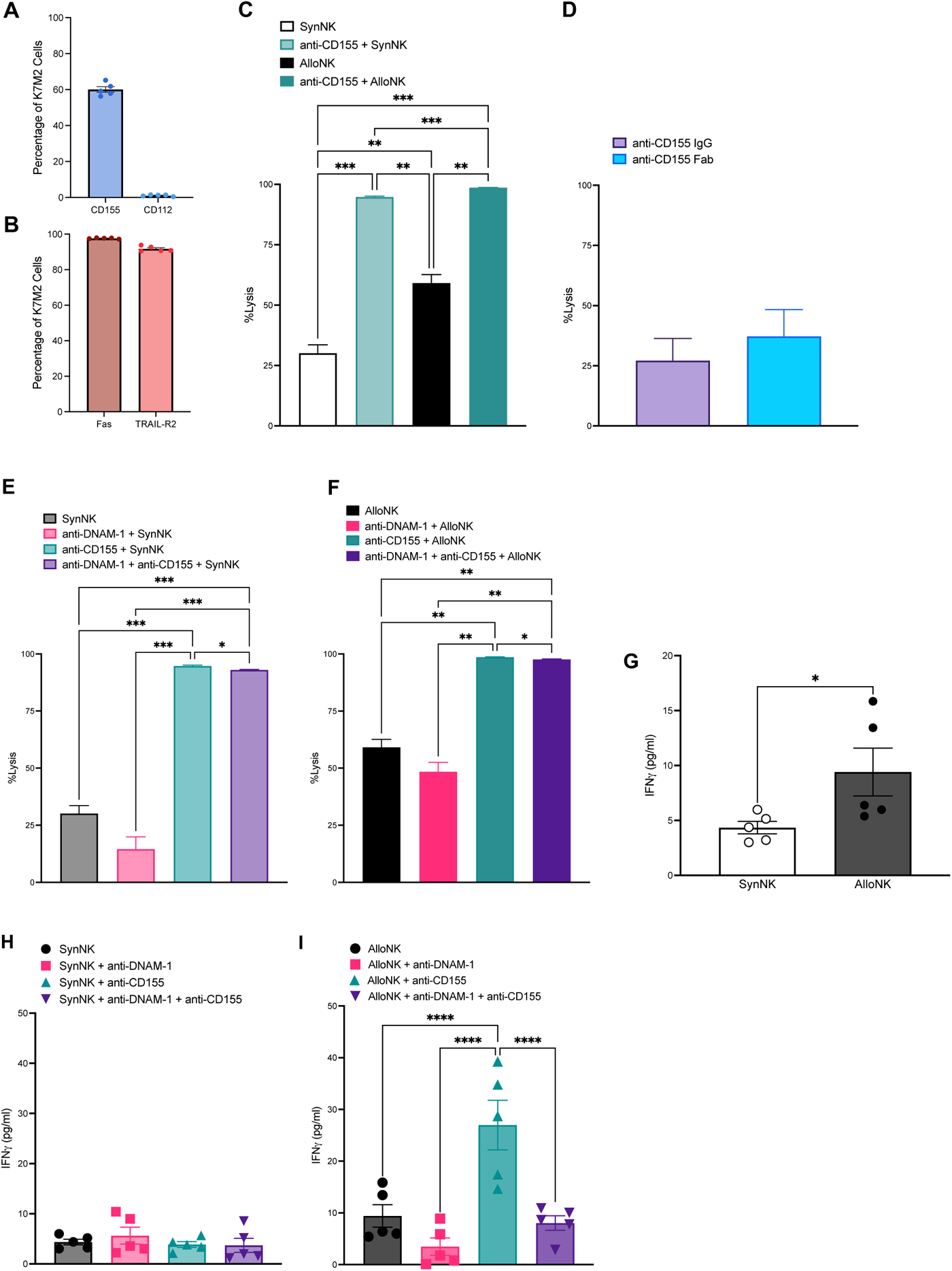
AlloNKs exhibit potent anti-OS activity enhanced by CD155 blockade. (A) Proportion of K7M2 OS expressing CD155 and CD112 shown (B) Proportion of K7M2 OS expressing Fas and TRAIL R2 shown. Individual values (n = 5) plotted with mean and SEM. EAE murine alloNK and synNK expanded for 12 days were plated with mKate2-expressing K7M2 OS at an effector: target ratio of 2.5:1 with or without anti-CD155 antibody in the presence of green fluorescent caspase 3/7-substrate. Green and red fluorescence objects were measured by Incucyte real-time analysis. %Lysis was calculated based on target alone and staurosporine-induced max lysis conditions. OS lysis shown for (C) alloNK and synNK with or without anti-CD155 antibody blockade at E:T 2.5:1 (n = 5 per group), (D) alloNK with anti-CD155 antibody or anti-CD155 Fab at E:T 2:1 (n = 3 per group), (E) synNK with anti-CD155, anti-DNAM-1, or anti-CD155 and anti-DNAM-1 blockade at E:T 2.5:1 (n = 5 per group), and (F) alloNK with anti-CD155, anti-DNAM-1, or anti-CD155 and anti-DNAM-1 blockade at E:T 2.5:1 (n = 5 per group). Supernatant was collected and analyzed for IFNγ release by ELISA with samples in triplicate. Concentration was calculated based on 5 parameter curve fit interpolated from standard curve. IFNγ concentration shown for (G) alloNK and synNK (H) synNK with anti-CD155, anti-DNAM-1, or anti-CD155 and anti-DNAM-1 blockade, and (I) alloNK with anti-CD155, anti-DNAM-1, or anti-CD155 and anti-DNAM-1 blockade. Mean with SEM shown for technical replicates. Two-sided two-sample t tests and one-way ANOVA and Kruskal-Wallis with Dunn’s multiple comparisons tests were performed (* p < 0.05, ** p < 0.01, *** p < 0.001, **** p < 0.0001).

Blocking CD155 could either enhance cytotoxicity by blocking engagement of TIGIT or CD96 on NK cells or impair cytotoxicity by inhibiting engagement with DNAM-1 on NK cells. It is also unclear if CD155 blockade can enhance cytotoxicity further with Ly49 mismatch. With CD155 blockade, both alloNK and synNK showed increased OS lysis after 24 hours (Fig. 2C). Notably, alloNKs with CD155 blockade yielded the highest OS lysis (98.60 ± 0.11%) and was significantly higher than OS lysis by synNKs with CD155 blockade (94.74 ± 0.37%) (Fig. 2C). Since NK cells express CD16, it is possible that blocking CD155 may inadvertently label the tumor for antibody-dependent cellular cytotoxicity (ADCC). To distinguish if alloNKs were engaging in ADCC, an antibody fragment (Fab) was generated containing only the antigen binding portion of the anti-CD155 antibody without the constant region necessary to induce ADCC via CD16 on NK cells. Compared to the full anti-CD155 antibody, the anti-CD155 Fab did not significantly affect alloNK OS lysis, confirming that the CD155 blocking effect was not mediated by ADCC (Fig. 2D). Additionally, a CD155-expressing human OS cell line, MG63, was similarly more sensitive to human NK-92 cell mediated tumor killing with CD155 blockade, suggesting these findings are translatable to other cell lines and species (Supplementary Fig 4 A-C).

Since DNAM-1 is an activating receptor that binds CD155, we determined whether DNAM-1 was necessary for NK cell cytotoxicity against CD155-expressing OS. Blocking DNAM-1 did not significantly affect OS lysis by alloNKs or synNKs (Fig. 2E-F), likely because NK cells have other activating receptors to recognize tumors. However, when DNAM-1 blockade was combined with CD155 blockade, OS lysis by alloNKs (97.63 ± 0.21%) and synNKs (93.01 ± 0.23%) was significantly decreased compared to CD155 blockade alone, though cytotoxicity remained markedly elevated compared to no blockade or DNAM-1 blockade alone (Fig. 2E-F).

We next evaluated how co-culture with CD155-expressing K7M2 OS affected cytokine production by murine NK cells. AlloNKs produced significantly more IFNγ (9.41 ± 2.18 pg/ml) compared to synNKs (4.35 ± 1.26 pg/ml) (Fig. 2G). While synNK IFNγ production was not affected by DNAM-1 blockade, CD155 blockade, or combination DNAM-1 and CD155 blockade, alloNK IFNγ production was significantly increased with CD155 blockade (26.97 ± 4.79 pg/ml) compared to no blockade (Fig. 2H-I). AlloNKs with DNAM-1 blockade alone or combination DNAM-1 and CD155 blockade had significantly lower IFNγ production compared to CD155 blockade alone (Fig. 2I), implying the DNAM-1/CD155 axis is a more potent regulator of IFNγ production than Ly49 mismatch alloreactivity.

Next, we investigated if CD155 or DNAM-1 blockade altered NK cell cytotoxicity mechanisms. First, degranulation was assessed by measuring CD107a surface expression on synNKs and alloNKs after incubation with K7M2 OS. AlloNKs had greater CD107a MFI and proportion of CD107a^+^ cells compared to synNKs after incubation with K7M2 OS (Fig. 3A). With the addition of CD155 blockade, DNAM-1 blockade, or combination CD155 and DNAM-1 blockade, there were no significant differences in CD107a MFI or the proportion of CD107a^+^ synNKs or alloNKs (Fig. 3B-C). AlloNKs also had increased perforin MFI and proportion of perforin^+^ cells compared to synNKs, but no significant differences in granzyme B MFI or %granzyme B^+^ cells (Fig. 3D). Overall, the proportion of FasL^+^ and TRAIL^+^ cells were low in synNKs and alloNKs, with alloNKs having greater proportions of these cells compared to synNKs (Fig. 3E). FasL MFI was significantly lower in alloNKs compared to synNKs. There were no changes in CD107a MFI, CD107a^+^, FasL MFI, Granzyme B MFI, FasL^+^, or Granzyme B^+^ in synNKs or alloNKs with CD155 axis blockade (Supplementary Fig. 3).

**Figure 3:**
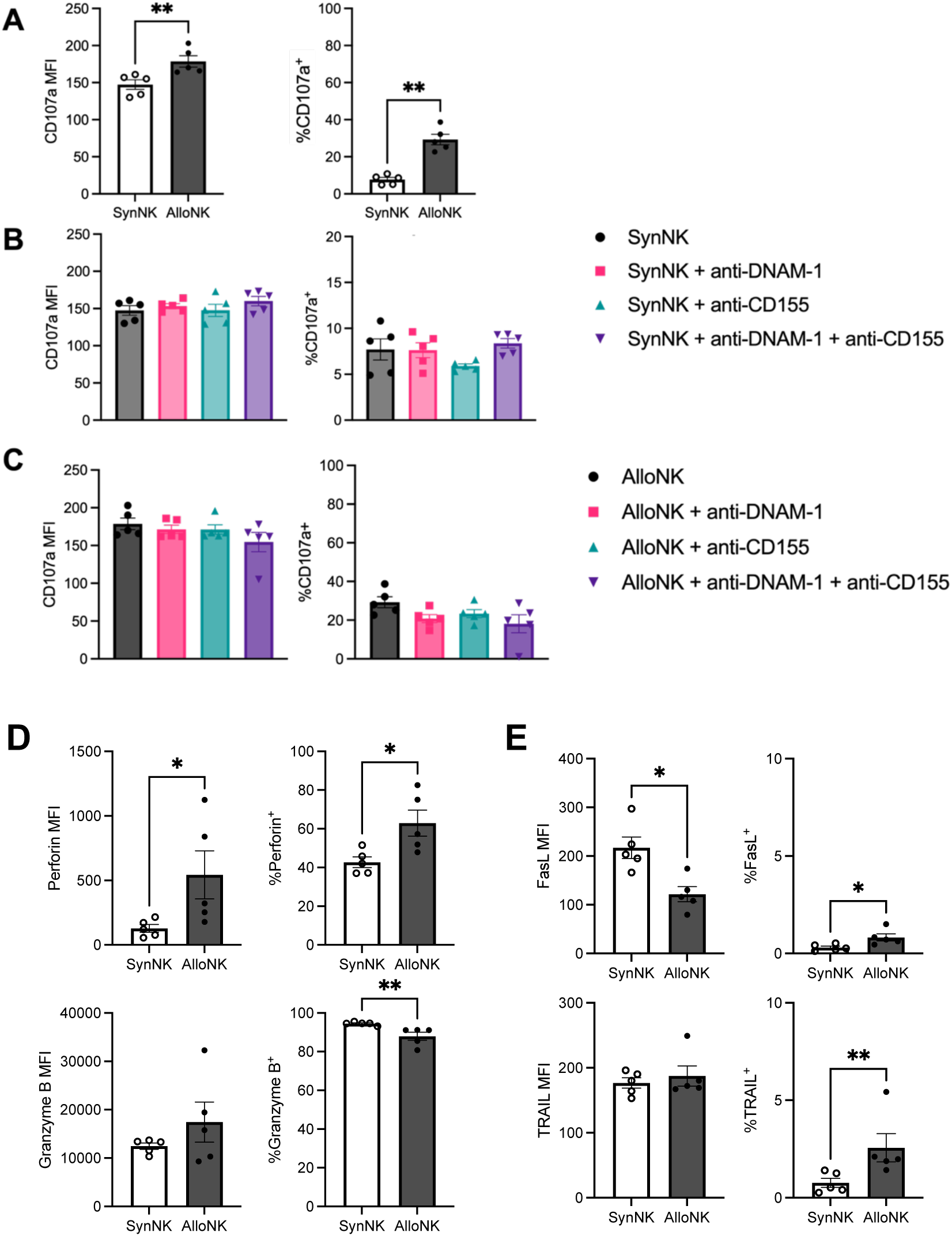
Degranulation is the main killing mechanism used by alloNKs against OS. EAE murine alloNK and synNK cells were plated with K7M2 OS at an E:T ratio of 2.5:1. Cells were incubated for 30 minutes, followed by addition of monesin and brefeldin A, and incubated for an additional 4 hours. NK cells were collected, fixed and permeabilized, stained with surface antibodies and analyzed by flow cytometry. Cells were gated on B6 (NK1.1^+^) and BALB/c (CD49b^+^) cells and CD107a MFI was calculated and gated on CD107a^+^ cells perforin^+^, granzyme B^+^, FasL^+^, and TRAIL^+^ cells. CD107a MFI and percentage of cells expressing CD107a for alloNK and synNK cells (A), synNK cells with CD155 axis blockade (B), and alloNK cells with CD155 axis blockade (C) are shown. MFI and percentage of NK cells expressing perforin, and granzyme B (D) and FasL and TRAIL (E) are shown. Mean with SEM shown for experimental replicates (n = 5). Two-tailed two-sample t-tests and one-way ANOVA with Kruskal-Wallis with Dunn’s multiple comparisons tests were performed (* p < 0.05, ** p < 0.01).

### AlloNK treatment with CD155 blockade increases survival after alloBMT in relapsed OS

Clinically, patients are typically offered allogeneic bone marrow transplant (alloBMT) when in remission or without active disease. Yet many patients will relapse after alloBMT. Due to superior activity observed in vitro, we investigated whether alloNK infusion with CD155 blockade could control recurrence of pulmonary OS metastases that occurred after alloBMT. To test this, BALB/c recipient mice underwent lethal irradiation and alloBMT with T-cell-depleted B6 bone marrow since T cells compete with NK cells for cytokines like IL-15 and can cause GVHD, limiting GVT effects. Seven days post-transplant, mice were inoculated with K7M2 OS intravenously (Fig. 4A) to mimic relapse of pulmonary metastases post-alloBMT. Mice were then treated with a single EAE alloNK cell infusion at day 14 given with either isotype antibody or anti-CD155 blockade.

**Figure 4:**
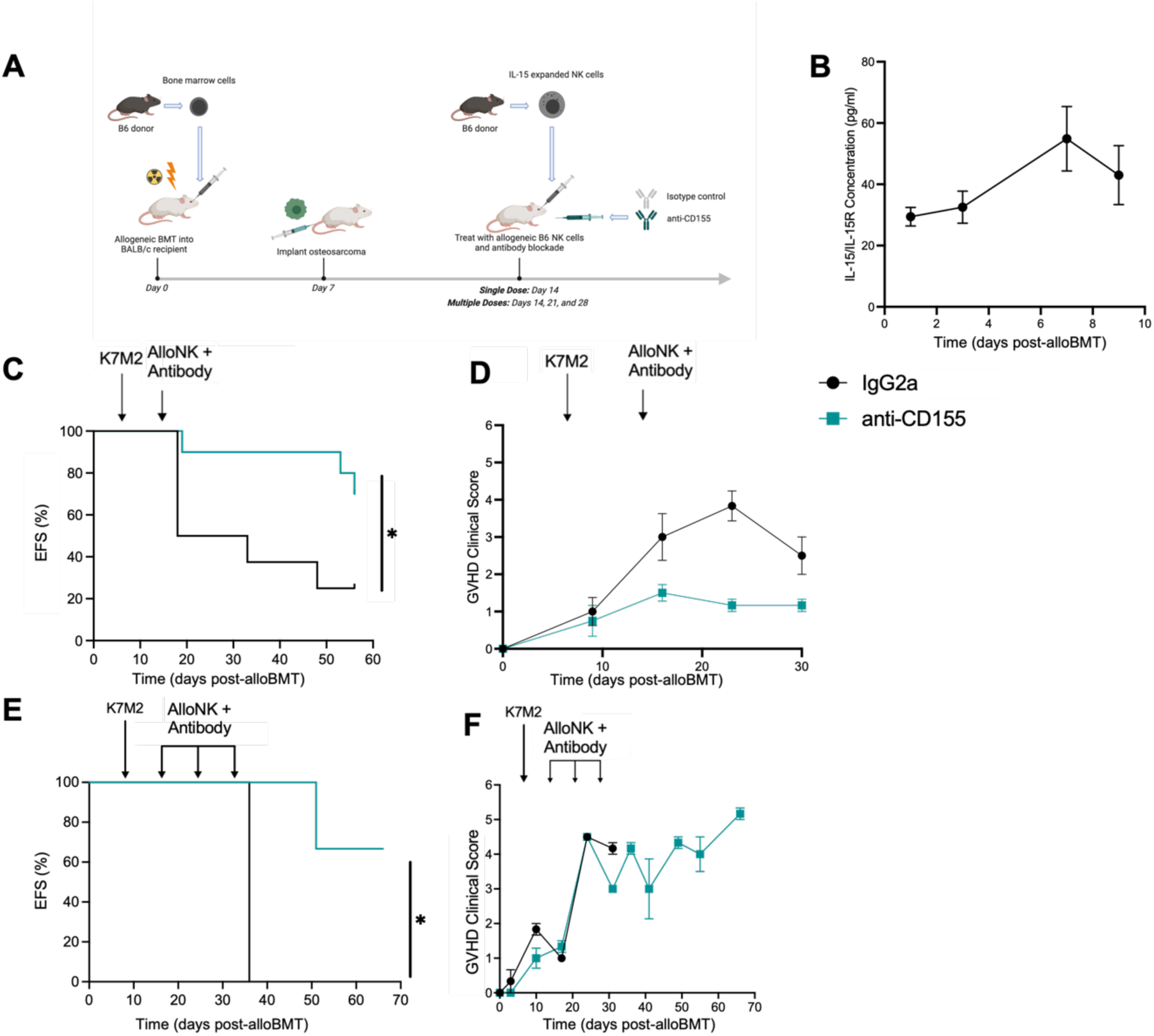
AlloBMT and single or multiple dose CD155 blockade with alloNK effectively treats relapsed OS pulmonary metastases. (A) Schematic showing alloBMT followed by K7M2 OS inoculation and single dose or multiple doses of alloNK infusion with CD155 axis blocking antibodies. (B) Serum levels of mouse IL-15/IL-15Rl1l pre- and post-BMT and single dose alloNK and CD155 axis blockade treatment at timepoints post alloBMT (n = 6-15 mice per timepoint) (C) EFS for mice treated with alloBMT and single dose alloNK with isotype control or anti-CD155 blockade are shown at timepoints post alloBMT (n = 8 or 10 mice per group). (D) Clinical GVHD scores for mice treated with alloBMT and single dose alloNK with isotype control or anti-CD155 blockade are shown (n = 8 or 10 mice per group). (E) EFS for mice treated with alloBMT and multiple doses of alloNK with isotype control or anti-CD155 blockade are shown at timepoints post alloBMT (n = 3 mice per group). (F) Clinical GVHD scores for mice treated with alloBMT and multiple doses of alloNK with isotype control or anti-CD155 blockade are shown (n = 3 mice per group). Bars and points with error bars depict mean with SEM. Log-rank test performed and student’s t-tests were performed (* p < 0.05).

Post-alloBMT, serum IL-15 increased (Fig. 4B). Treatment with alloNK and anti-CD155 blockade significantly increased event-free survival (EFS), with an event defined as tumor burden increased by at least four-fold or subject death, compared to alloNK with isotype control (Fig. 4C). EAE NK cell infusions after alloBMT have been associated with exacerbating GVHD in humans (28). There was no significant difference in GVHD between mice treated an EAE NK cell infusion with isotype antibody or anti-CD155 blockade (Fig. 4D). Thus, anti-CD155 antibody does not exacerbate GVHD from residual T cells in the donor graft, graft-derived de novo T cells or EAE alloNK infusion. Treatment with multiple doses of EAE alloNK cell infusions with anti-CD155 blockade resulted in similar improvement in EFS without exacerbation of GVHD (Fig. 4E, F).

### Combination alloBMT, alloNK treatment and CD155 blockade does not control established pulmonary OS disease

Because there are limited options for patients with relapsed/refractory OS, we then tested the efficacy of using T-cell-depleted alloBMT followed by EAE alloNK cells and anti-CD155 blockade to treat mice with established pulmonary OS metastases. BALB/c mice were inoculated with K7M2 OS intravenously to introduce experimental metastases (Fig. 5A). Mice were then treated with an alloBMT using T-cell-depleted B6 bone marrow cells at day 7, followed by EAE alloNK cells infusion and isotype control or anti-CD155 blocking antibody on day 8. To determine the role of DNAM-1 on the in vivo effects of CD155 blockade, EAE alloNK cells pre-incubated with anti-DNAM-1 blocking antibody were infused with either an isotype antibody or anti-CD155 blockade. Post-alloBMT, again serum levels of IL-15 increased (Fig. 5B). In an established disease setting, there was no significant EFS benefit in mice treated with alloBMT followed by anti-CD155 blockade compared to mice treated with alloNKs and isotype antibody (Fig. 5C). When alloNKs were infused in the setting of anti-DNAM-1 blockade, there was no significant difference in EFS when compared to mice receiving isotype control or anti-CD155 blockade (Fig. 5C). Anti-CD155 or anti-DNAM-1 blockade in this model did not exacerbate GVHD (Fig. 5D). Tumor burden was not significantly different between mice treated with anti-CD155 blockade or isotype antibody (Fig. 5E). To determine persistence of infused alloNK cells, EAE alloNK expressing CD45.1 were measured by flow cytometry at 6, 13, and 16 days after infusion and were detectable up to 16 days after infusion (Fig. 5F). Loss of target antigen can mediate immune escape despite presence of effector immune cells. Examination of tumor-bearing lung tissue post-tumor inoculation confirmed marked tumor burden and persistent CD155 expression *in vivo* (Supplementary Fig. 5). Unlike treating relapsed disease after alloBMT, treatment of established pulmonary OS metastases with alloBMT followed by alloNK cells and CD155 blockade did not significantly impact disease progression.

**Figure 5:**
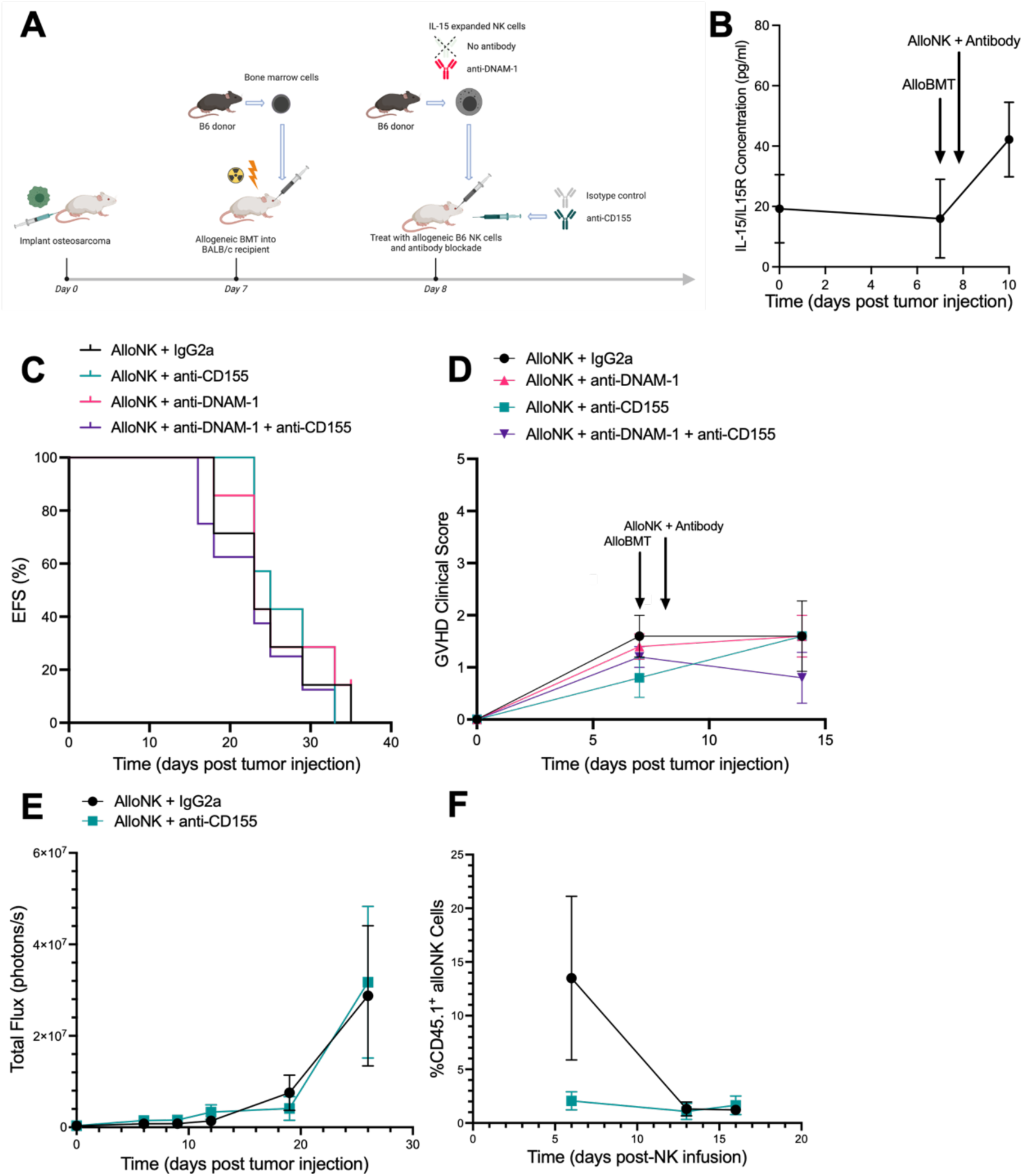
CD155 blockade with alloNK shows limited efficacy against established OS disease after alloBMT. (A) Schematic showing K7M2 OS inoculation, alloBMT, and alloNK infusion with CD155 axis blocking antibodies. (B) Blood sera levels of mouse IL-15/IL-15Rl1l pre- and post-BMT and alloNK and CD155 axis blockade treatment (n = 7-8 mice per timepoint) (C) EFS for mice treated with alloBMT and alloNK with isotype control, anti-DNAM-1 blockade, anti-CD155 blockade, or anti-DNAM and anti-CD155 blockade are shown (n = 7-8 mice per group). (D) Clinical GVHD scores for mice treated with alloBMT and alloNK with isotype control, anti-DNAM-1 blockade, anti-CD155 blockade, or anti-DNAM and anti-CD155 blockade are shown (n = 7-8 mice per group). (E) Tumor burden measured by IVIS shown as total flux (photons/s) for mice treated with alloBMT and alloNK with isotype control or anti-CD155 blockade (n = 6 per group) (F) Recovery of infused alloNK expressing CD45.1 from spleens harvested from mice treated with alloBMT and alloNK with isotype control or anti-CD155 blockade (n = 3 mice per timepoint). Bars and points with error bars depict mean with SEM.

### Effects of combination alloBMT, alloNK and CD155 axis blockade in vivo on gene expression of OS lung metastases

To investigate changes in the TME within the established pulmonary OS metastases model that may explain resistance to alloBMT and alloNK therapy, we performed RNA microarray analysis on lung tissue isolated from OS tumor-bearing mice treated with alloNK with or without anti-DNAM-1 pre-treatment and either IgG2a isotype or anti-CD155 antibody. Focusing on differential gene expression changes of the CD155 blockade treatment compared to isotype control, among genes with an absolute fold change greater than or equal to 1.5, 3 genes were upregulated, and 5 genes were downregulated (Fig. 6B). Genes associated with increased NK cytotoxicity and immune cell infiltration were upregulated in CD155-blockade treated mice (Fig. 6B). Ccl3, encoding chemokine ligand 3, was upregulated in the CD155 blockade treated mice with a 1.790-fold increase (adjusted p value = 0.038). Granzyme A (*Gzma*) had a 1.729-fold change (adjusted p value = 0.038) whereas Chemokine ligand 9 (*Ccl9)* had a 1.558-fold change (adjusted p value = 0.038). In addition, *Vegfa* was downregulated with a fold change of -.0624 (adjusted p-value = 0.03). *Vegfa* encodes vascular endothelial growth factor A, which is regulated by immune cells and associated with tumor vascularization (29). *Lamp3*, or lysosomal-associated membrane protein 3, was found to be downregulated in lung tissue from CD155 blockade treated mice with a fold change of 0.547 (adjusted p-value = 0.011). Dimensional analysis showed that gene changes in the CD155 blockade treated group were orthogonal to changes in the group treated with anti-DNAM-1 pretreated EAE NKs (Fig. 6D).

**Figure 6:**
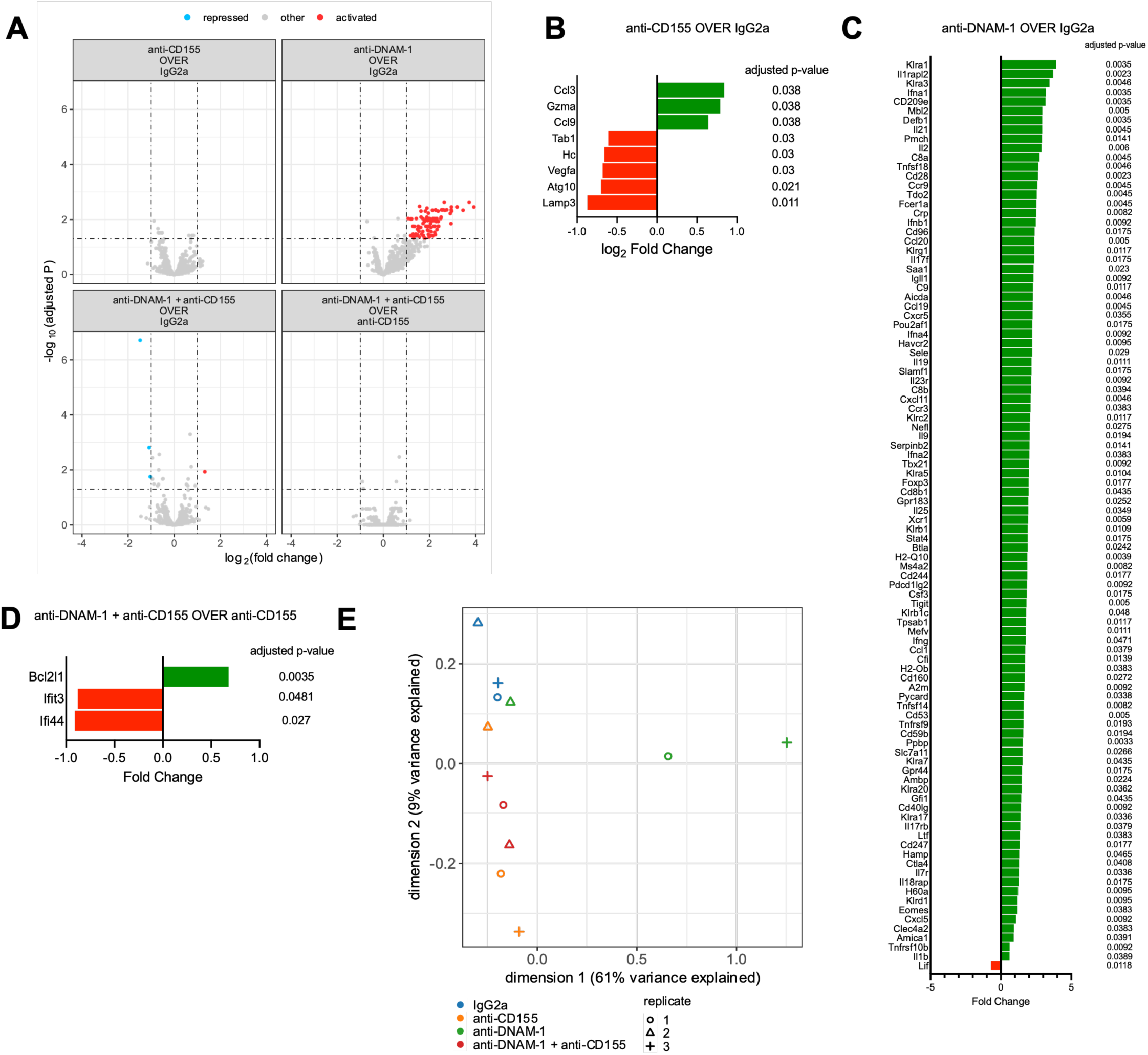
CD155 blockade and DNAM-1 blockade with alloNK treatment changes gene expression in lung tissue of OS tumor-bearing mice post alloBMT. (A) Volcano plots of differential expression shown for treatment groups. A horizontal cutoff is drawn at adjusted P = 0.05. Two vertical cutoffs are drawn at log2(fold change) at -1 and 1. (n = 3 experimental replicates per group). Differentially expressed genes with absolute log2(fold change) ≥ 0.6 are plotted with adjusted p value ≤ 0.05 are shown for (B) mice treated with alloNK and anti-CD155 over alloNK and IgG2a (C) mice treated with alloNK and anti-DNAM-1 over alloNK and IgG2a and (D) mice treated with alloNK, anti-DNAM-1, and anti-CD155 over alloNK and anti-CD155. E) Multidimensional scaling distances between endogenous genes quantified from each NanoString dataset shown. Distance between samples on this plot corresponds to log-fold-change on gene expressions between samples (n = 3 experimental replicates per group).

In mice treated with anti-DNAM-1 pretreated EAE NKs, genes related to NK cell function were upregulated relative to mice treated with isotype control (Fig. 6C). Various Ly49 receptors were found to be upregulated in the mice treated with anti-DNAM-1 pretreated EAE NKs including *Klra1* (Ly49A), *Klra3* (Ly49C), Klrg1 (CLEC15A), Klrc2 (NKG2C), Klra5 (Ly49E), Klrb1 (CD161), Klrb1c (NK1.1), Klra7 (Ly49G), Klra20 (Ly49T), Klra17 (Ly49Q), and Klrd1 (CD94) (Fig. 6C). *H60a*, which encodes the NKG2D ligand H60, was also found to be upregulated after DNAM-1 blockade (Fig. 6C). Other genes encoding inhibitory immune cell receptors were upregulated in mice treated with anti-DNAM-1 pretreated EAE NKs, including Cd96 (CD96), tigit (TIGIT), ctla4 (CTLA-4) and havcr2 (TIM-3) (Fig. 6C).

Changes in differential gene expression in mice treated with anti-DNAM-1 pretreated EAE NKs indicated a role for modulating T cell responses (Fig. 6C). Various tumor necrosis factor (TNF) superfamily members were upregulated in lung tissue of DNAM-1 treated mice. *Tnfsf18* was upregulated with a fold change of 2.66 (adjusted p value = 0.0046). *Tnfsf18* is known as GITR or activation inducible TNFR family member (AITR), a type I transmembrane protein that is expressed at high levels on CD4^+^CD25^+^ regulatory T cells (Tregs) and on B, NK, NKT cells, granulocytes and macrophages (30). *Tnfrs9,* encoding TNFR family member 9, was significantly upregulated in mice treated with DNAM-1 pretreated EAE NKs with a fold change of 1.61 (adjusted p value = 0.0193).

Comparing mice treated with anti-CD155 blockade and alloNK cells pre-treated with anti-DNAM-1 blockade to mice receiving anti-CD155 blockade and alloNK cells, only 1 gene was upregulated, and 2 genes were downregulated (Fig. 6D). Examining multidimensional scaling distance between the combination of anti-CD155 with anti-DNAM-1 blockade and anti-CD155 blockade alone, similar changes occurred (Fig. 6E). Thus, additional DNAM-1 blockade on EAE NKs did not impact differential gene expression with CD155 blockade.

### DNAM-1 blockade has more potent inhibitory effects compared to NKG2D blockade

Due to differential gene expression change of the NKG2D ligand *H60a* after anti-DNAM-1 EAE NKs infusion *in vivo*, we investigated how DNAM-1 and NKG2D blockade compare in modulating alloNK cell function. Murine synNK and alloNK cells were co-cultured with K7M2 OS with DNAM-1 blocking antibody and evaluated for NKG2D expression. AlloNK cells had higher NKG2D MFI and proportion of NKG2D^+^ NK cells compared to synNK cells (Fig. 7A). SynNKs had markedly increased NKG2D expression and proportion of NKG2D^+^ NK cells when co-cultured with K7M2 OS with DNAM-1 and CD155 blockade combined compared to isotype control, and the combination of DNAM-1 and CD155 blockade significantly increased NKG2D^+^ NK cells compared to CD155 blockade alone (Fig. 7B). In alloNKs, DNAM-1 blockade alone increased NKG2D MFI and NKG2D^+^ NK cells compared to isotype control and CD155 blockade alone (Fig. 7B). To see if NKG2D upregulation augmented cytotoxicity, alloNK cells were tested for cytotoxicity against K7M2 OS with anti-DNAM-1, anti-NKG2D, and anti-CD155 blockade. While DNAM-1 blockade significantly decreased tumor lysis when combined with anti-NKG2D, NKG2D blockade did not significantly change tumor lysis when added to CD155 blockade (Fig. 7C). These data indicate that DNAM-1 is a more potent regulator of alloNK cell anti-OS responses compared to NKG2D.

**Figure 7:**
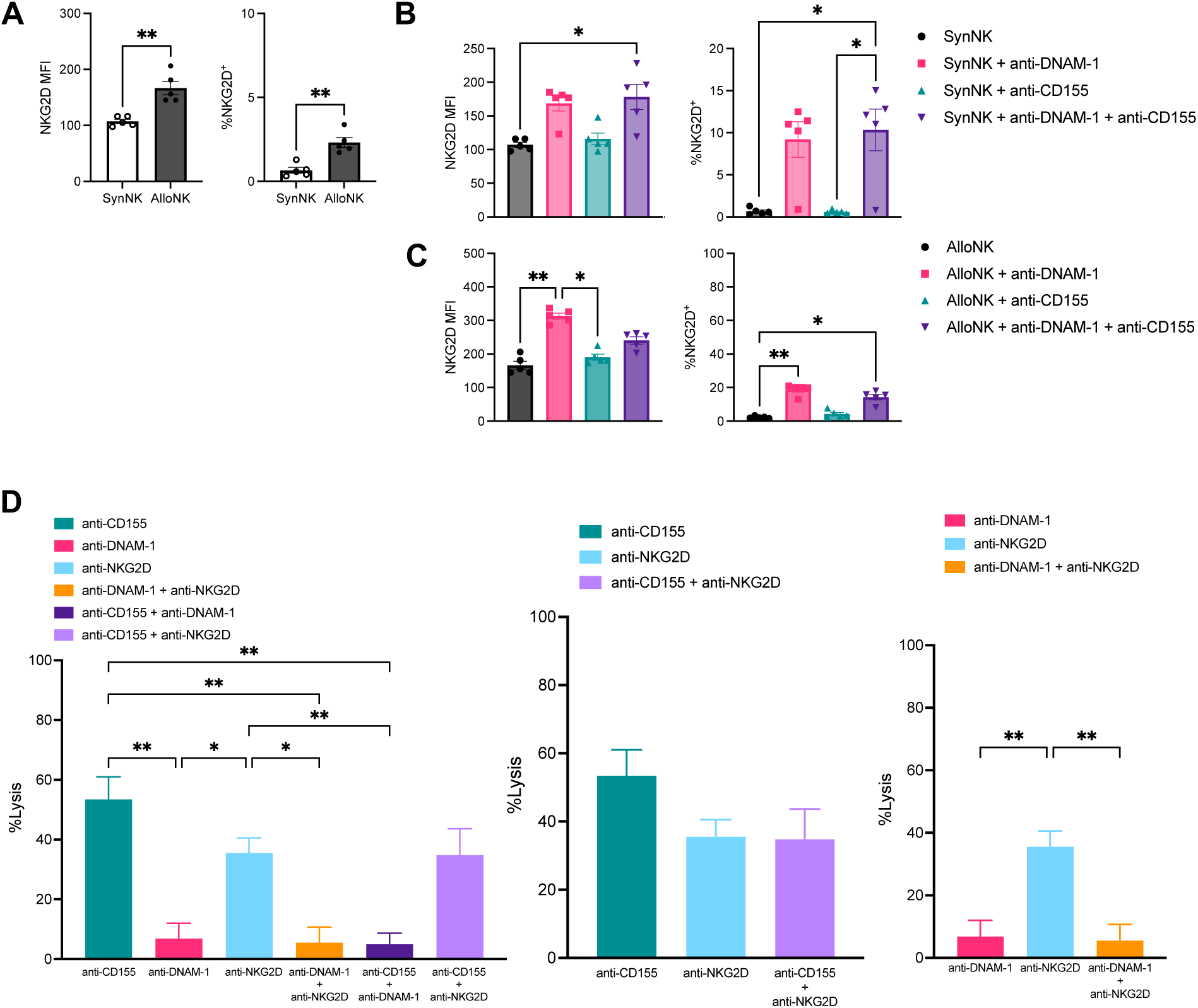
DNAM-1 blockade increases NKG2D expression in synNK and alloNK cells and has a greater impact on cytotoxicity compared to NKG2D. EAE murine alloNK and synNK were plated with K7M2 OS at an E:T ratio of 2.5:1. NK cells were collected and analyzed by flow cytometry. NKG2D MFI and percentage of NKG2D^+^ NK cells shown for (A) alloNK and synNK (B) synNK with anti-CD155, anti-DNAM-1, or anti-CD155 and anti-DNAM-1 blockade, and (C) alloNK with anti-CD155, anti-DNAM-1, or anti-CD155 and anti-DNAM-1 blockade. Mean with SEM shown for experimental replicates (n = 5). EAE murine alloNK and synNK were plated with mKate2-expressing K7M2 OS. %Lysis was calculated based on target alone and staurosporine-induced max lysis conditions. OS lysis shown for (D) alloNK and synNK with combinations of anti-CD155, anti-DNAM-1, and anti-NKG2D blockade at E: T 2.5:1 (n = 5 per group). Two-tailed two-sample t-tests and one-way ANOVA with Kruskal-Wallis with Dunn’s multiple comparisons tests were performed (* p < 0.05, ** p < 0.01).

## DISCUSSION

NK cells have been studied for decades to treat advanced solid tumors in adults and children, but with inconsistent success. The emergence of more refined clinical alloBMT regimens that minimize GVHD (e.g. α/β T cell receptor depletion, post-transplant cyclophosphamide, etc) provides a platform to incorporate adoptive cell therapy with alloNK cells and enhance GVT effect against solid tumors, but understanding the relative contributions of both NK cell activating and inhibitory receptors to GVT effects in this environment is needed.

In this study, murine alloNKs demonstrated greater degranulation and increased perforin expression against OS compared to synNKs, suggesting that Ly49 receptor mismatch impacts NK cell-mediated cytotoxicity. With respect to translatability, human immunoglobulin-like receptors (KIRs) and murine Ly49 receptor exhibit polymorphic and polygenic heterogeneity and serve analogous functions (31). Activating receptor ligands differ by species as human NKG2D binds MICA, MICB, and ULBP family members, but murine NKG2D binds Rae-1 family members, MULT1, and H60 isoforms (32). Freshly isolated murine NK cells exhibit weaker lytic activity compared to freshly isolated human NK cells, depending on culture and stimulation conditions (33). Therefore, murine NK cell findings must be validated in human preclinical studies. We demonstrated that a CD155-expressing human OS cell line was more susceptible to NK-cell-mediated lysis with CD155 blockade (Supplementary Fig 4A-C). These preliminary data show translational potential, but further work is needed with xenograft and humanized models.

Granule exocytosis was impaired in synNKs, likely due to self-MHC recognition on OS. While FasL and TRAIL expression are increased in activated, mature NK cells, few EAE synNK or alloNK cells expressed these ligands after co-culture with CD155-expressing OS. While CD155 blockade enhanced NK cell cytotoxicity and IFNγ production against OS, combination CD155 and DNAM-1 blockade did not affect degranulation or death receptor expression in NK cells or affect IFNγ production by synNK cells. These results demonstrate that NK cell inhibition through self-MHC recognition regulates receptor expression independent of cytokine production. Blocking DNAM-1 does not significantly affect OS lysis by synNKs, as observed by others, (23), and we observed similar findings on alloNK cytotoxicity, IFNγ production, and degranulation. The K7M2 OS cell line has low expression of CD112 (Fig. 3A), which also interacts with DNAM-1 and TIGIT. Therefore, our model was limited in assessing CD112 and its impact on alloreactivity.

Blocking CD155 could either interfere with engagement of the activating receptor DNAM-1 on NK cells, leading to larger tumors, or engagement of the inhibitory receptors TIGIT or CD96 on NK cells, leading to smaller tumors. We evaluated CD155 blockade combined with alloNK infusion after alloBMT to treat two scenarios of pulmonary OS. First, we treated mice with pulmonary OS disease induced after alloBMT to mimic post-transplant relapse. These results showed that combining anti-CD155 antibody and alloNKs improves EFS and controls tumor growth in relapsed disease, perhaps interfering with TIGIT or CD96 inactivation of NK cells. Importantly, CD155 blockade or infusion of anti-DNAM-1 pretreated alloNKs did not exacerbate GVHD. A TIGIT-Fc fusion protein that binds to CD155 has been shown to activate signaling on DCs and reduce acute GVHD (34). To our knowledge, this is the first evidence that blocking CD155 after alloBMT is feasible and safe when combined with alloNK infusion.

While alloBMT with adoptively transferred alloNK and CD155 blockade improved survival from OS relapse after alloBMT, it was unsuccessful in treating established disease existent before alloBMT. To better understand barriers to success, we explored gene expression profiling of lung tissue in mice with established pulmonary OS treated with alloBMT, alloNKs, and CD155 blockade. Increased gene expression of *Gzma*, *Ccl9*, and *Ccl3* was observed in mice treated with CD155 blockade, consistent with immune cell effector activity and mobilization. Considering the role of DNAM-1, an activating CD155 receptor, various genes for NK inhibitory receptors, like Ly49A, Ly49C, Ly49E, CLEC15A, Ly49G, Ly49Q, and CD94 were upregulated in mice treated with anti-DNAM-1 EAE alloNKs. *Klrd1* (encodes CD94) forms a heterodimer with NKG2A, an inhibitory receptor encoded by *klrc1* that recognizes the non-classical class I molecule HLA-E in humans and Qa-1b in mice (35). NKG2A/CD94 engagement with HLA-E causes ITIM phosphorylation and SHP-1 recruitment to suppress effector functions (36). Thus, infusing anti-DNAM-1 EAE alloNKs caused upregulation of inhibitory Ly49 molecules and CD94 and ultimately no GVT effect. Strategies that block Ly49 inhibitory receptors and/or maximize DNAM-1 stimulation may skew alloNK cells to maximal GVT activity.

While NKG2D synergizes with other activating receptors like 2B4, it does not synergize with DNAM-1 (37). Thus, when alloNKs pre-treated with anti-DNAM-1 are infused, NKG2D may be available to stimulate NK cell antitumor effects. Increased gene expression of *H60a*, which encodes a potent NKG2D ligand, was observed in the pulmonary OS tissue. Interestingly, H60a recruits NK cells and sensitizes tumors to rapid NK cell-mediated elimination (38). Thus, in tumors expressing NKG2D ligands, any antitumor response impaired through attenuated DNAM-1 could theoretically be rescued by engaging the NKG2D pathway. However, blocking NKG2D did not significantly abrogate alloNK cell-mediated anti-OS cytotoxicity, unlike DNAM-1 blockade (Fig. 7D). Thus, in this context, DNAM-1 is a more potent regulator of alloNK cell cytotoxicity of OS compared to NKG2D. Further studies into regulation and hierarchy of the DNAM-1 and NKG2D pathways in alloNK cells may elucidate their relative activity with checkpoint blockade.

In conclusion, we found evidence that CD155 blockade with adoptively transferred alloNK cells significantly enhances the GVT effect against CD155-expressing pulmonary OS that relapses after alloBMT. When using alloBMT with alloNK infusions and anti-CD155 as a platform for treating established pulmonary OS, the GVT effect falters. Blocking DNAM-1 on alloNK cells increases inhibitory receptors that may prevent optimal GVT activity, even if NKG2D ligands are present, implying a hierarchy of receptors drives alloNK cell control of OS after alloBMT. These data are timely given that clinical trials incorporating α/β T cell depletion alloBMT as a means of depleting GVHD-causing α/β T cells while enriching the donor graft with GVT-promoting γ/δ T cells and NK cells are underway, including for children with OS (<u>NCT02508038</u>). A pilot trial testing EAE alloNK infusions in non-transplanted OS patients is also underway (<u>NCT03209869</u>) (39). These findings demonstrate that rebuilding the immune system after alloBMT using adoptive cell therapy and CD155 axis modulation is a potential approach for controlling relapsed, metastatic OS that occurs after alloBMT.

## List of Abbreviations

OS: osteosarcoma
EFS: event-free survival
NK: natural killer
KIR: killer-immunoglobulin-like receptor
MHC: major histocompatibility complex
HSCT: hematopoietic stem cell transplant
GVHD: graft-versus-host disease
GVT: graft-versus-tumor
EAE: ex vivo activation and expansion
TME: tumor microenvironment
E: T: effector: target
alloBMT: allogeneic bone marrow transplant
IFNγ: interferon gamma
MFI: median fluorescence intensity
TNF: tumor necrosis factor
synNK: syngeneic natural killer
alloNK: allogeneic natural killer
ADCC: antibody-dependent cellular cytotoxicity
Fab: antibody fragment

## Declarations

### Availability of data and material

The datasets used and/or analyzed during the current study are available from the corresponding author on reasonable request.

### Authors’ contributions

MMC was responsible for experimental design, execution, and analysis of data, created final versions of all figures and drafted the manuscript. LSh isolated RNA and performed quality control for microarray analysis. AEG transduced tumor cell lines with lentiviral vector. LSo, AEQ, FS, JE, DT, DB, JK, EL, MFP, and AC contributed to experimental design and data collection. CMC provided experimental design, review of data, and revised the manuscript. All authors read and approved the final manuscript.

## Acknowledgements

We thank the Division of Hematology, Oncology and Bone Marrow Transplant in the Department of Pediatrics and Carbone Cancer Center at the University of Wisconsin-Madison for their ongoing support. The author(s) thank the UWCCC Flow Laboratory, Small Animal Imaging and Radiotherapy Facility, Experimental Animal Pathology lab, Translational Research Initiatives in Pathology, UWCCC Informatics Shared Resource and Biomedical Research Model Services for use of their facilities and services.

