## Supplemental Figures for "CD155 blockade enhances allogeneic natural killer cell-mediated antitumor response against osteosarcoma"

### Supplementary Figure 1.

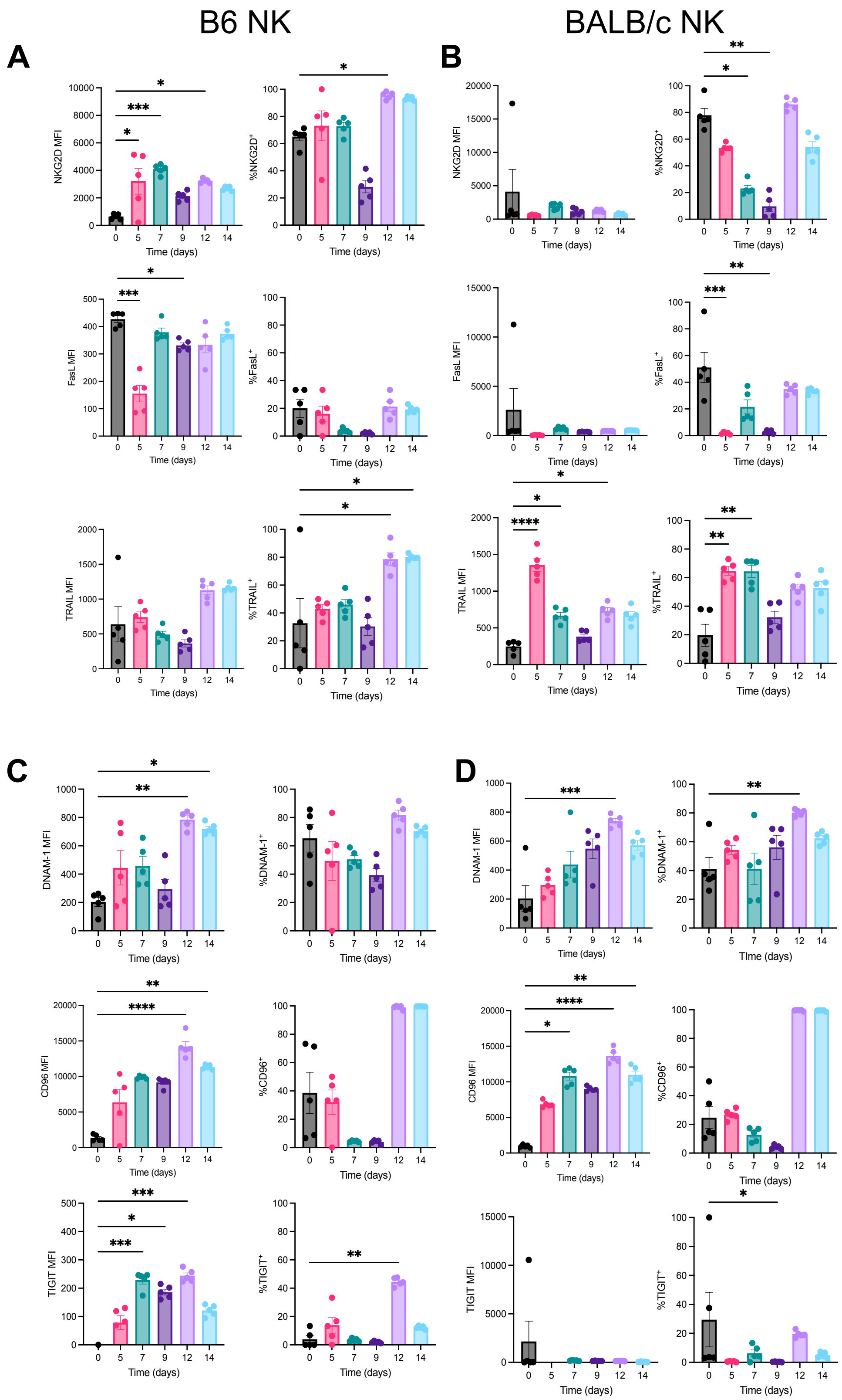

**Supplementary Figure 2.**

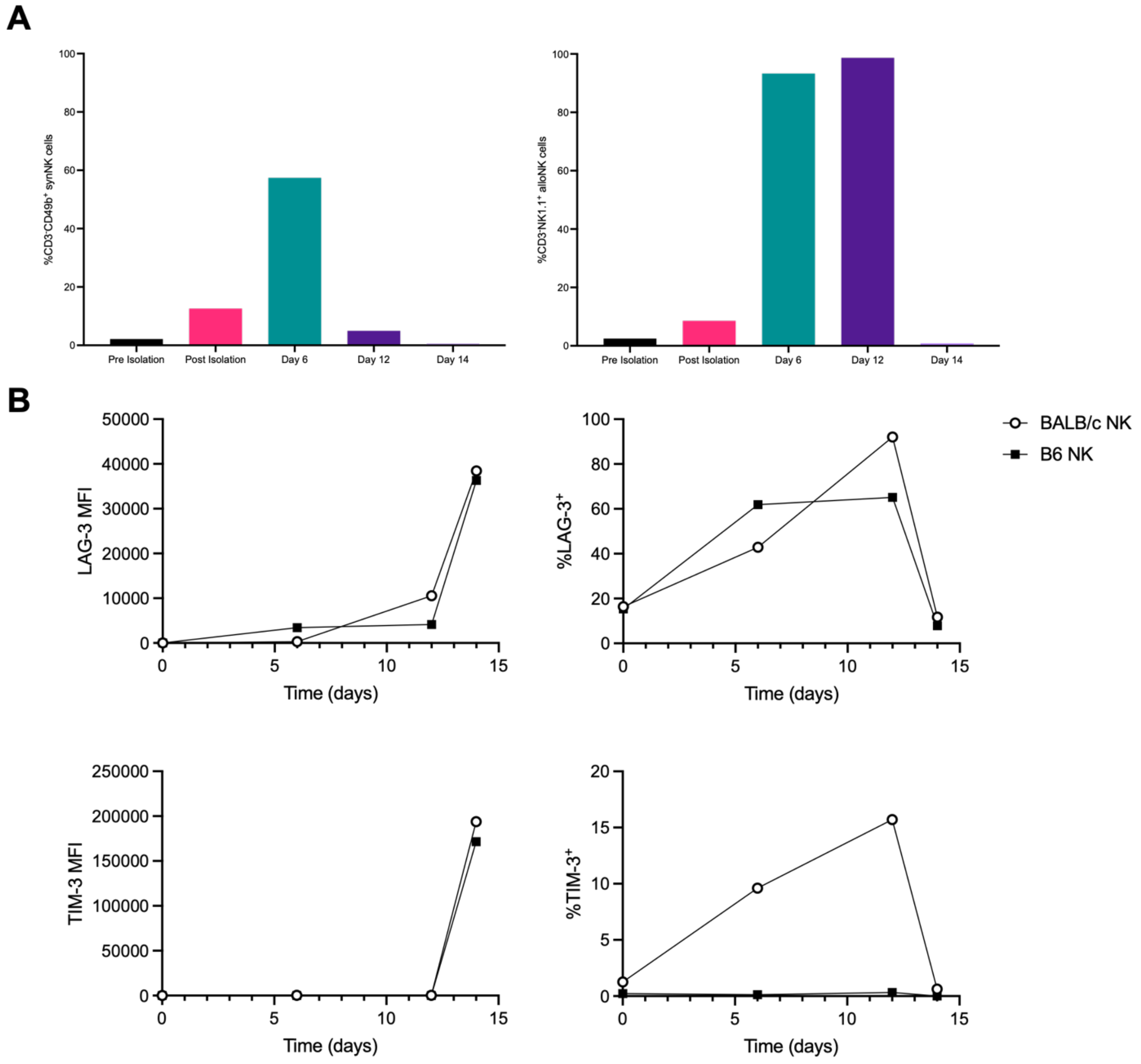

### Supplementary Figure 3.

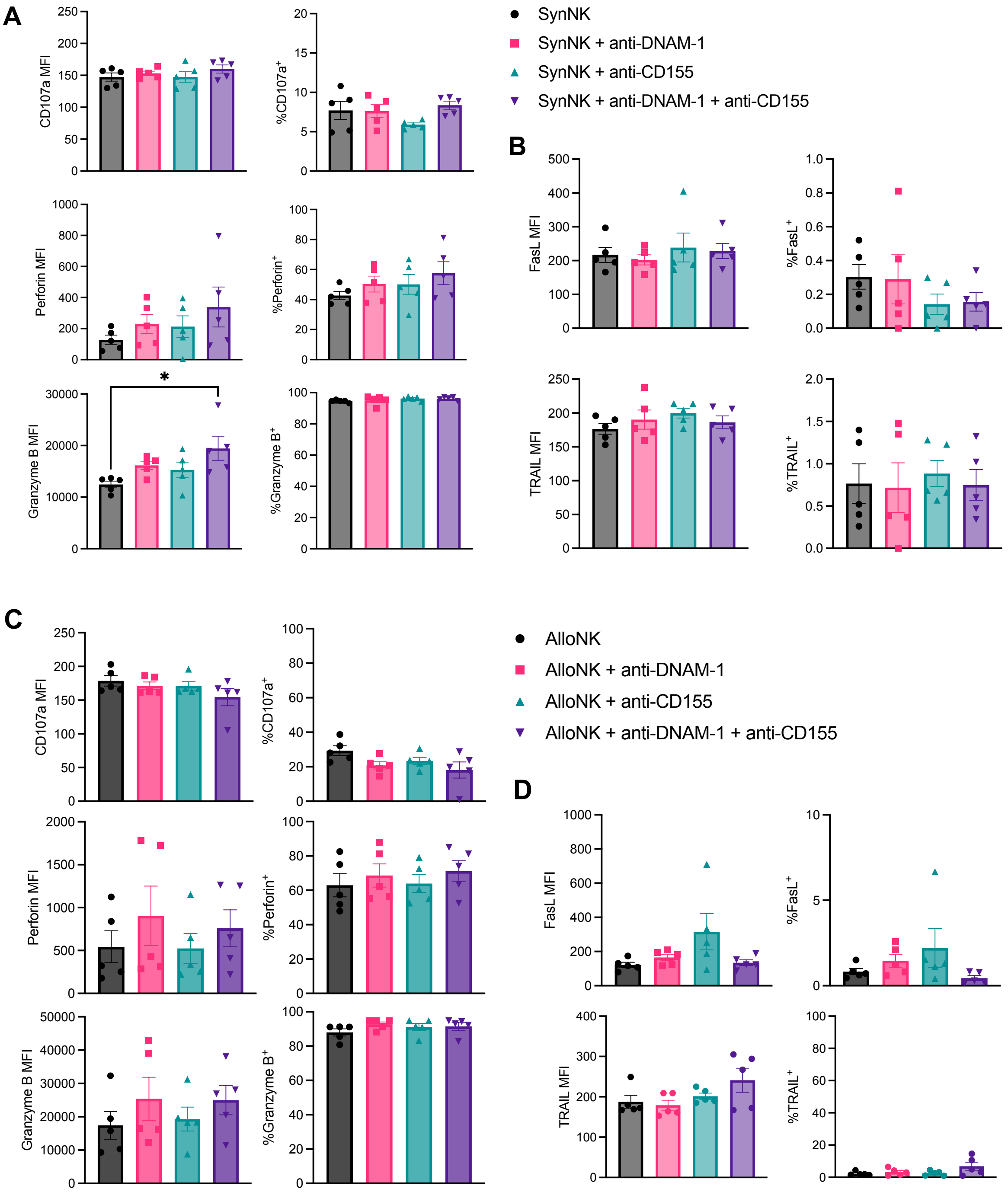

### Supplementary Figure 4.

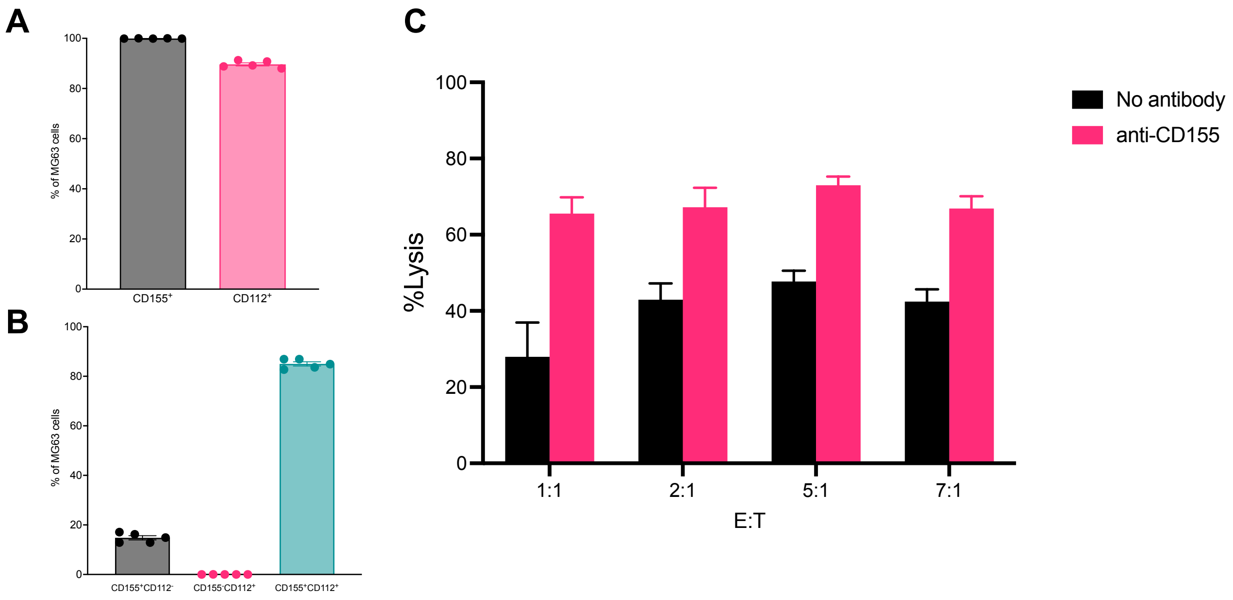

**Supplementary Figure 5.**

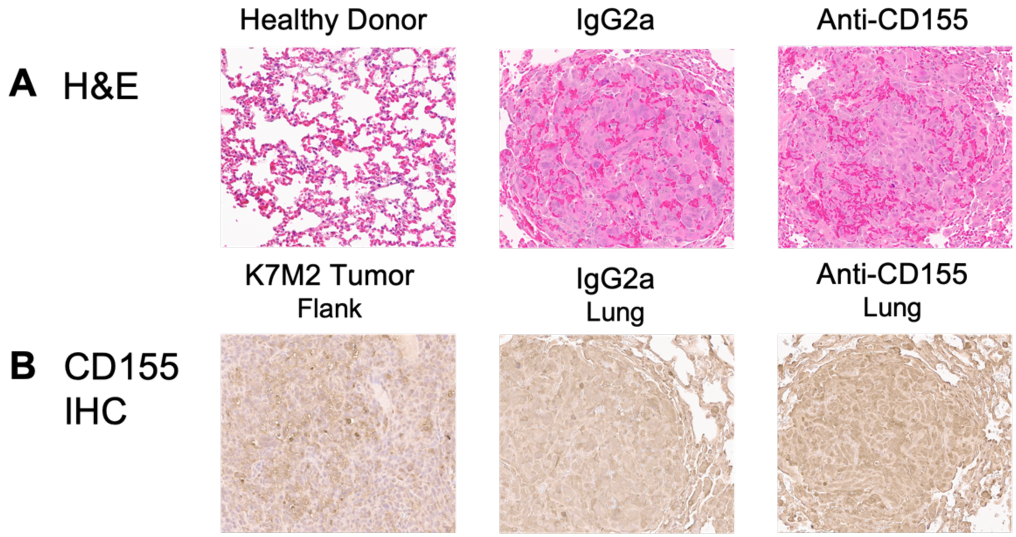
