## Supplemental Figure Captions for "CD155 blockade enhances allogeneic natural killer cell-mediated antitumor response against osteosarcoma"

**Supplementary Figure 1: EAE murine NK cells have increased expression of NKG2D, TNF death receptors, and CD155 receptors DNAM-1, CD96 and TIGIT.** B6 (NK1.1+) and BALB/c ($CD49b^+$) NK cells were harvested and analyzed for expression of activating receptor DNAM-1 and inhibitory receptors CD96 and TIGIT by flow cytometry on days 0, 5, 7, 9, 12, and 14. DNAM-1 MFI and %DNAM-$1^+$, CD96 MFI and %$CD96^+$, and TIGIT MFI and %$TIGIT^+$ of (A) B6 and (B) BALB/c NK cells are shown. DNAM-1 MFI and %DNAM-$1^+$, CD96 MFI and %$CD96^+$, and TIGIT MFI and %$TIGIT^+$ of (C) B6 and (D) BALB/c NK cells are shown.

**Supplementary Figure 2: EAE NK cell product has high purity at day 12 and increased exhaustion at day 14.** (A) Purity of B6 and BALB/c NK cells shown pre- and post-isolation and on days 6, 12, and 14 of expansion. (B) Expression of exhaustion markers LAG-3 (LAG-3 MFI, %LAG-$3^+$) and TIM-3 (TIM-3 MFI, %TIM-$3^+$) on BALB/c and B6 NK cells on days 6, 12, and 14 of expansion are shown.

**Supplementary Fig. 3: Degranulation is not significantly affected by CD155 or DNAM-1 blockade in synNK or alloNK cells against OS.** EAE murine alloNK cells plated with K7M2 OS at an E:T ratio of 2.5:1. Cells were incubated for 30 minutes, followed by addition of monesin and brefeldin A, and incubated for an additional 4 hours. NK cells were collected, fixed and permeabilized, stained with surface and intracellular antibodies and analyzed by flow cytometry. Cells were gated on B6 ($NK1.1^+$) cells and CD107a, perforin, granzyme B, FasL, and TRAIL MFI were calculated. NK cells were further gated on $CD107a^+$, $perforin^+$, granzyme $B^+$, $FasL^+$, and $TRAIL^+$ cells. MFI and percentage of synNK cells expressing CD107a, perforin, and granzyme B (A) and FasL and TRAIL (B) and alloNK cells expressing CD107a, perforin, and granzyme B (C) and FasL and TRAIL (D). Mean with SEM shown for experimental replicates (n = 5). One-way ANOVA with Kruskal-Wallis with Dunn's multiple comparisons tests were performed.

**Supplementary Fig. 4 CD155 blockade increases human NK-92 cell-mediated lysis of CD155-expressing MG63 OS cell line.** (A) Proportions of MG63 OS expressing CD155 and CD112 and (B) proportions of MG63 OS expressing only CD155, only CD112 or both CD155 and CD112 are shown. NK-92 cells were plated at a 1:1, 2:1, 5:1 and 7:1 E:T ratios with MG63 $GFP^+$ cells with or without anti-CD155 antibody. Mean and SEM shown for n=3 replicates per condition.

**Supplementary Fig. 5. CD155 expression in pulmonary tumor disease is persistent despite anti-CD155 blockade therapy in vivo** (A) H&E stain of lung tissue samples from a healthy donor BALB/c mouse and a BALB/c mice inoculated with K7M2 tumor intravenously after treatment with alloBMT on day 7 and alloNK with isotype control or anti-CD155 blockade on day 8. Samples taken day 13 after tumor inoculation. (B) Samples obtained from a subcutaneously implanted K7M2 right flank tumor from a BALB/c mouse after 16 days of tumor growth and lung tissue samples from BALB/c mice inoculated with K7M2 tumor intravenously after tumor injection on day 13 after treatment with alloBMT on day 7 and alloNK with isotype control or anti-CD155 blockade on day 8 were stained for CD155 expression by IHC. All images shown at 20X magnification.
