## Supplemental Methods for "CD155 blockade enhances allogeneic natural killer cell-mediated antitumor response against osteosarcoma"

### Supplementary Methods

#### Cell lines

Murine K7M2 OS cells (ATCC, Manassas, VA) were cultured in DMEM medium with 10% fetal bovine serum with 50 IU/mL penicillin/streptomycin, 1X MEM nonessential amino acids, 25 mM HEPES buffer, and 1 mM sodium pyruvate. Human MG-63 OS cells (ATCC) were cultured in DMEM with 10% FBS and 50 IU/mL penicillin/streptomycin. Human NK-92 cells (ATCC) were cultured in RPMI with 10% FBS and 100 IU/ml rhIL-2 (Biologic Resources Branch, National Cancer Institute (NCI)). Cell authentication was performed using short tandem repeat analysis (Idexx BioAnalytics, Westbrook, ME) and per ATCC guidelines using morphology, growth curves, and *Mycoplasma* testing within 6 months of use with MycoStrip Mycoplasma Detection Kit (Invitrogen, Waltham, MA). Cell lines were maintained in culture at 37°C in 5% $CO_2$. K7M2 were transduced to express the luminescent reporter luciferase or the fluorescent reporter mKate2 using a lentiviral vector pLV[Exp]-Puro-EFS>SV40 NLS/mKate2 (VectorBuilder, Chicago, IL).

#### Generation of anti-CD155 Fab fragment

Purified rat anti-mouse anti-CD155 antibody (Leinco Cat. No. C2833, Fenton, MO) was digested using the Pierce Fab preparation kit (Pierce Biotechnology Cat. No. 44985, Rockford, IL). To verify Fab product, Fab product and undigested IgG were analyzed under reducing conditions by SDS-Page on a 10% Bis-Tris gel ((source?) to verify molecular weights.

#### Flow cytometric analysis

Tumors and spleens were removed, mechanically dissociated, and passed through a 70μm cell strainer (Corning, Glendale, AZ). Cell suspensions were centrifuged at 300g for 10 minutes, and then digested with ACK lysing buffer. Cells were then washed and centrifuged at 300g for 10 minutes and resuspended in 10 mL PBS and counted on the COULTER COUNTER® Z1 Series Particle Counter (Beckman Coulter, Brea, CA).  Then, $1x10^6$ cells were added to flow cytometry tubes in staining buffer (phosphate-buffered solution with 2% FBS) and stained with DNAM-1 FITC (Biolegend Cat. No. 128803), CD253 (TRAIL) PerCP-Cy5.5 (Biolegend Cat. No. 142108), CD96 APC (Biolegend Cat. No. 131712), CD49b Pacific Blue (Biolegend Cat. No. 109918), NK1.1 BV510 (Biolegend Cat. No. 108738) FasL PE (Biolegend Cat. No. 106606), TIGIT PE-Cy7 (Biolegend Cat. No. 142108), and GhostRed780 viability dye (Tonbo Biosciences Cat. No. 13-0865-T100).

For intracellular cytokines, cells were fixed using BD Cytofix buffer and permeabilized with BD Perm/Wash buffer. Cells were stained with CD107a-1 FITC (Biolegend Cat. No. 121606), CD253 (TRAIL) PerCP-Cy5.5 (Biolegend Cat. No. 142108), FasL APC (Biolegend Cat. No. 106610), Granzyme B AF700 (Biolegend Cat. No. 372222), CD49b Pacific Blue (Biolegend Cat. No. 109918), NK1.1 BV510 (Biolegend Cat. No. 108738), Perforin PE (Biolegend Cat. No. 154306), NKG2D PE-Dazzle (Biolegend Cat. No. 131712), LAG-3 PE (Biolegend Cat. No. 125208), TIM-3 APC (Biolegend Cat. No. 134008), CD45.1 APC (Biolegend Cat. No. 110714) and GhostRed780 viability dye (Tonbo Biosciences Cat. No. 13-0865-T100). K7M2 OS were stained with CD155 PE (Biolegend Cat. No. 337610), CD112 (R&D Systems, Cat. No. FAB3869G), TRAIL-R2 PE (Biolegend Cat. No. 119906), and Fas APC (Biolegend Cat. No. 152604).

Cells were analyzed using the Attune NXT flow cytometer (Thermo Fisher, Waltham, MA). Flow data were analyzed using FlowJo 10 (FlowJo, Ashland, OR). Cells were gated on forward scatter and side scatter singlets and live cells. B6 NK cells were defined as NK1.1$^{+}$ cells and BALB/c NK cells were defined as CD49b$^{+}$ cells. NK cell populations were further defined by NKG2D$^{+/-}$, DNAM-1$^{+/-}$, CD253 (TRAIL) $^{+/-}$, CD96 $^{+/-}$, FasL $^{+/-}$, and TIGIT $^{+/-}$, and median fluorescence intensity was measured for NKG2D, DNAM-1, CD253 (TRAIL), CD96, and TIGIT.

**NK cytotoxicity assay**

EAE B6 and BALB/c murine NK cells were plated with murine K7M2 mKate2 OS in 96 well flat bottom plates at effector to target (E: T) ratio of 2:1 or 2.5:1, with target cell seeding density at 10,000 cells/well and effector cell density at 20,000 or 25,000 cells/well, with n = 3 or 5 wells per condition. For antibody blockade, cells were incubated with purified rat anti-mouse anti-CD155 antibody (Leinco Cat. No. C2833) or purified rat anti-mouse anti-DNAM-1 (Biolegend Cat. No. 128822) antibody for 30 minutes at 37°C prior to use. As a maximum killing positive control, target K7M2 mKate2 was incubated with 1 uM staurosporine. As a negative control, cells were incubated with 10 ug/mL rat IgG2a antibody (Leinco Cat. No. I-1177). For detection of apoptosis by caspase-3/7 activity, 5 uM NucView488 Caspase-3 (Biotium Fremont, CA) was added to each well according to manufacturer's protocol. Plates were imaged using the IncuCyte live imaging system (Sartorius, Goettingen, Germany) at 10X magnification with images taken every 4 hours for 24 hours. Cell killing by target cell lysis was measured by evaluating the number of target cells present undergoing apoptosis.

**Murine NK cell degranulation and cytokine production assays**

EAE B6 and BALB/c murine NK cells were plated with murine K7M2 mKate2 osteosarcoma in 96 well flat bottom plates at effector to target (E: T) ratio of 2.5:1, with target cell seeding density at 100,000 cells/well and effect cell density at 250,000 cells/well, with n = 5 wells per condition. For antibody blockade, cells were incubated with 10 ug/ml rat IgG2a antibody (Leinco Cat. No. I-1177), purified rat anti-mouse anti-CD155 antibody (Leinco Cat. No. C2833), or purified rat anti-mouse anti-DNAM-1 (Biolegend Cat. No. 128822) antibody for 30 minutes at 37°C prior to use in assay. For degranulation and other flow cytometry analysis, anti-CD107a FITC antibody was added to plated cells. After incubation for 30 minutes at 37°C, GolgiSTOP (monesin) and GolgiPLUG (brefeldin A) (source?) were added to each well, and the cells were incubated for another 4 hours at 37°C. Cells were harvested for flow cytometry analysis. For IFNγ production analysis, plated cells were incubated for 4 hours at 37°C and supernatant was collected and frozen at -80°C for later analysis by ELISA.

**IFNγ ELISA**

Collected undiluted cell supernatant was assayed for IFNγ detection using Mouse IFNγ ELISA MAX (Biolegend Cat. No. 430815). Peripheral blood samples were collected from the submandibular vein using a lancet in a serum separation tube, and serum was collected and saved at -80°C. Undiluted sera were assayed using a Mouse IL-15/IL15R ELISA kit (Invitrogen Cat. No. BMS6023, Waltham, MA). ELISA plates were read using a CLARIOstar microplate reader (BMG Labtech, Cary, NC) at 450 nm wavelength. Analyte concentration was determined based on a non-linear 5-parameter logistic curve calculated using Prism 9 (Graphpad Software, San Diego, CA).

**Subcutaneous K7M2 tumor**

To generate an in vivo control K7M2 tumor, a BALB/c mouse was inoculated with 1E6 subcutaneous luciferase$^+$ K7M2 murine OS cells in the right flank. After 16 days, the mouse was euthanized and the tumor excised.

**Immunohistochemistry**

Murine tissues were excised and fixed in 10% buffered formalin for 24 to 48 hour, then transferred to 70% ethanol until processed for paraffin embedding and cut into 5 μm sections. Sections were stained with hematoxylin and eosin. For IHC, sections were deparaffinized with xylenes and hydrated through graded alcohols to water. Antigen retrieval was performed in citrate buffer pH 6.0 (10 mM citric acid, 0.05% tween 20) for 3 minutes in the Biocare Decloaker.  Endogenous peroxidase was blocked with 0.4% $H_2O_2$ in phosphate-buffered saline for 10 minutes at room temperature. Slides were then serum blocked with 10% goat serum (Sigma) for 1 hour at room temperature, followed by primary rabbit anti-human anti-CD155 antibody (LS-VBIO, LS-B12331) at 1:200 dilution in 1% goat serum overnight at 4ºC. SignalStain IHC Detection (HRP, Rabbit) was then applied for 30 minutes at room temperature, washed with PBS, followed by DAB substrate (Cell Signaling Technology) and

Mayer's hematoxylin (Sigma). Slides were then dehydrated through graded alcohols to xylene. All slide images were scanned with a 20x objective using the Aperio Digital Pathology Slide Scanner.

**Infused alloNK recovery**

To evaluate infused alloNK recovery, EAE CD45.1-expressing alloNK cells were generated from B6-Ly5.1/Cr mice and infused into BALB/c recipient mice post tumor inoculation and alloBMT. At various timepoints, mice were euthanized and their spleens were harvested, processed into single-cell suspensions, and analyzed by flow cytometry to identify infused alloNK ($CD45.1^{+}CD3^{-}NK1.1^{+}$) and distinguish them from BMT-derived alloNK ($CD45.1^{-}CD3^{-}NK1.1^{+}$).

**nCounter RNA microarray and analysis**

Lung tissue was harvested from mice (n = 3 per group) post-euthanasia and processed into single cell suspensions. Total RNA was isolated from lung tissue from individual mice using a total RNA isolation kit (Qiagen, Germantown, MD, USA) according to the manufacturer's protocol. RNA underwent quality control checks for A260/230, A260/280 and DV200 readings and were analyzed using the nCounter Mouse PanCancer Immune Profiling Panel Kit (XT_PGX_MmV1_CancerImm_CSO) (NanoString Technologies, Inc., Seattle, WA, USA). Gene expression levels were quantified by the R package NACHO (version 2.0.4) from NanoString's RCC files. Differential gene expression was performed by the Bioconductor package edgeR (version 3.40.0). A differentially expressed gene was required to have at least two-fold changes and an adjusted $P < 0.05$. GSEA was carried out by the Bioconductor package fgsea (version 1.24.0) with nCounter PanCancer Immune Profiling Panel mouse gene sets downloaded from NanoString website. Raw microarray data have been deposited at the NCBI GEO under the accession number GSE252084.
